# Unravelling native plant resistomes – The *Sphagnum* microbiome harbours versatile and novel antimicrobial resistance genes

**DOI:** 10.1101/695973

**Authors:** Melanie-Maria Obermeier, Julian Taffner, Alessandro Bergna, Anja Poehlein, Tomislav Cernava, Christina Andrea Müller, Gabriele Berg

## Abstract

The expanding antibiotic resistance crisis calls for a more in depth understanding of the importance of antimicrobial resistance genes (ARGs) in pristine environments. We, therefore, studied the microbiota associated with *Sphagnum* forming the main vegetation in undomesticated, evolutionary old bog ecosystems. In our complementary analysis of a culture collection, metagenomic data and a fosmid library, we identified a low abundant but highly diverse pool of resistance determinants, which targets an unexpected broad range of antibiotics including natural and synthetic compounds. This derives both, from the extraordinarily high abundance of efflux pumps (80%), and the unexpectedly versatile set of ARGs underlying all major resistance mechanisms. The overall target spectrum of detected resistance determinants spans 21 antibiotic classes, whereby β-lactamases and vancomycin resistance appeared as the predominant resistances in all screenings. Multi-resistance was frequently observed among bacterial isolates, e.g. in *Serratia, Pandorea, Paraburkhotderia* and *Rouxiella.* In a search for novel ARGs we identified the new class A β-lactamase Mm3. The native *Sphagnum* resistome comprising a highly diversified and partially novel set of ARGs contributes to the bog ecosystem’s plasticity. Our results shed light onto the antibiotic resistance background of non-agricultural plants and highlight the ecological link between natural and clinically relevant resistomes.

---

The risk posed to modern medicine by increased morbidity and mortality associated with antibacterial resistance continues to escalate globally and has reached a stage where a post-antibiotic era is not unthinkable anymore^1,2^. Many of the clinically relevant antimicrobial resistance genes (ARGs) originate from the environment, wherein they may act in intra-community signalling and metabolic processes; in the presence of selective pressure they can adapt antibiotic resistance as primary function^3^. In order to retrace the origin and habitat transitions of resistant microorganisms, a detailed understanding of native resistomes is crucial^3^. So far, such elucidations focused on soil, water and air^4^. Limited work has been performed on plants and thereby mostly evolved around fresh produce to assess the risk potential of crops in serving as gateway of ARGs to humans^5–7^. The resistome of native plants from pristine vegetation was neglected so far. It can, however, provide the missing ecological link to understand the evolution and functioning of native resistomes as well as their role as pools of unexplored resistance mechanisms^8^. Since the resistome reflects the continuous co-evolution of small bioactive molecules and microbial genomes within an environment^9^, native plants, which provide an extraordinarily diversified secondary metabolism, are expected to possess a diversified intrinsic resistome as well.

*Sphagnum magellanicum* Brid. covering peatlands, was selected as a model plant to study ARGs in a representative pristine as well as evolutionary old ecosystem^10,11^. *Sphagnum*-dominated peatlands constitute balancing and productive ecosystems, in which the prevailing harsh conditions fostered symbiotic connections throughout a long plant-microbe co-evolution^12,13^. As a result, the *Sphagnum* microbiome is highly abundant and diverse with a specialised structure and function similar across geographic locations^14,15^. The microbiota fulfils important functions like nutrient supply and protection against biotic and abiotic stress; its metagenome is characterised to a remarkably high extend by signatures indicating horizontal gene transfer and communication systems thought to facilitate the balance between plasticity and stability within the bog ecosystem^13^. Moreover, the highly stable microbiome is not affected by soil microbiota, given that the rootless *Sphagnum* moss grows on peat; accumulated, partly degraded plant material mostly stemming from the plant itself and forming the largest terrestrial carbon sink on Earth^11^. *Sphagnum* mosses harbour specific and rich metabolite profiles^13^ and their associated microbiota is characterised by a high proportion of antimicrobial activity^16^. Altogether, *S. magellanicum* represents an ideal model to elucidate the antibiotic resistance background of plants which we expect to: i) comprise predominantly resistances against natural antibiotics due to the missing selective pressure by synthetic ones, ii) encompass versatile but evenly distributed ARGs due to the diverse and stable microbial community, iii) contain yet-unknown resistance genes. For our study, we pursued a unique approach combining analysis of a culture collection, *in silico* data mining of deep-sequenced metagenomic data, and functional metagenomics; the importance of combinatorial approaches for functional validation of *in silico* predictions was emphasised but rarely considered before^5,17^.

## RESULTS

### *Sphagnum* isolates display predominantly resistance against (semi)synthetic antibiotics

The culture collection was established using *Sphagnum* gametophytes from an Austrian Alpine bog, well known to host a highly abundant microbiota (Figure 1). Resistance assessment of the bacterial isolates included ten different antibiotics, comprising those ranked as critically important for medical applications^18^. Of the 264 isolated bacteria, 90% grew in the presence of at least one antibiotic, thereby, displaying 121 different resistance profiles (Supplementary Data 1). With predominantly observed vancomycin, ampicillin, rifampicin, ciprofloxacin, and sulfadiazine resistance, resistance against semisynthetic and synthetic antibiotics was more prevalent than resistance to natural antibiotics (Figure 2). This contradicts our preliminary expectation of dominating resistances against natural antibiotics. Overall, resistance against all ten antibiotics was observed. Multi-resistance against eight and nine antibiotics was encountered for three isolates each (Supplementary Data 1). The six isolates all displayed a distinct resistance profile, whereby all grew in the presence of erythromycin, vancomycin, rifampicin, ciprofloxacin and sulfadiazine. They were identified as the plant beneficial bacterium *Paraburkholderia phytofirmans*, formerly *Burkholderia phytofirmans*^19^, the potential nosocomial species *Serratia marcescens*, and as the newly described bacteria *Rouxiella chamberiensis, Pandoraea terrae* and *P. apista.*

**Figure 1:**
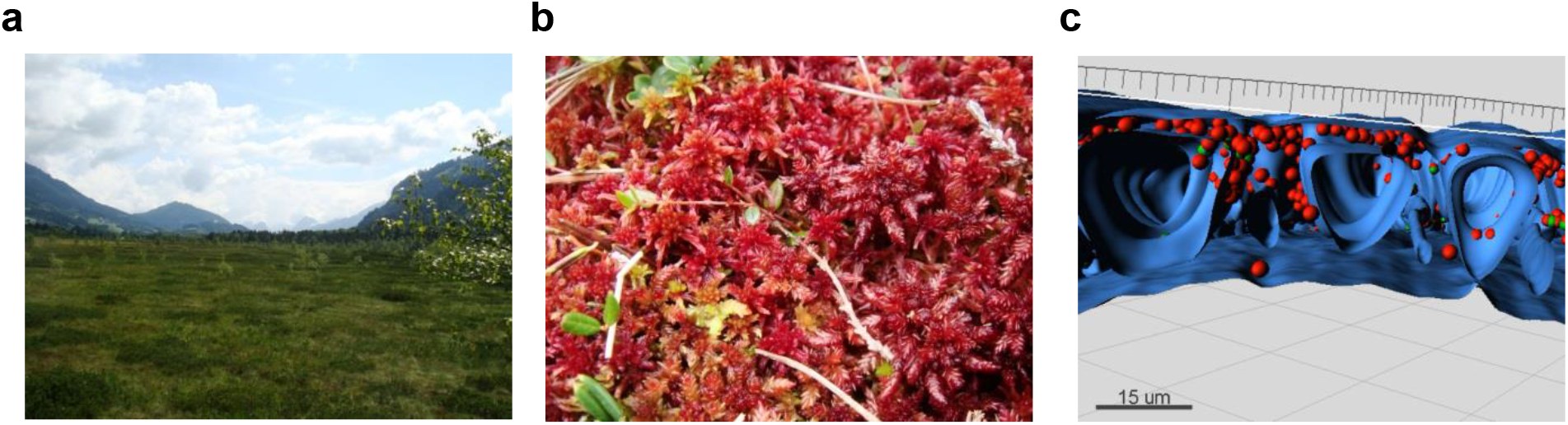
The *Sphagnum*-dominated peat bog. a) The Austrian Alpine peat bog Pürgschachen Moor (N47°34′50.57″ E14°20′29.29″). b) *S. magellanicum* gametophytes. c) Cross-section of a *Sphagnum* gametophyte displaying the highly abundant microbial colonisation (red spheres) of the moss’s hyaline cells (blue).

**Figure 2:**
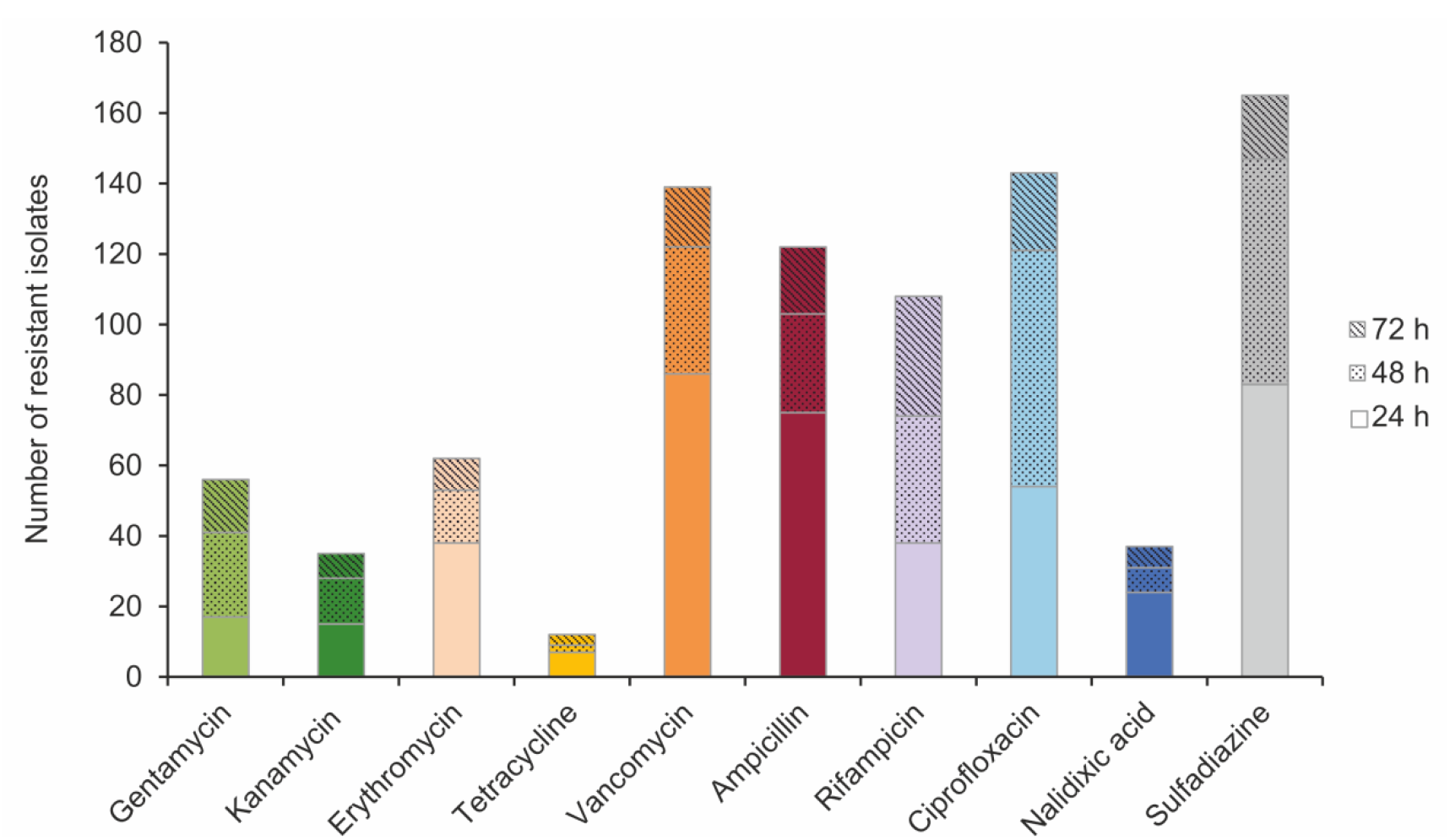
Antibiotic resistance profile of the *S. magellanicum* culture collection. Absolute number of resistant moss isolates which grew in the presence of different antibiotics.

### The *Sphagnum* microbiome comprises a highly diverse resistome

An lllumina-sequenced *S. magellanicum* metagenome was aligned against Comprehensive Antibiotic Resistance Database (CARD) sequences using high stringency (90% threshold) yielding matches with ≥30 amino acids identity. This revealed a low abundant, but highly diverse pool of resistance determinants (Supplementary Data 2). After curation and double normalisation of the generated hits (Supplementary Table 1), 0.14% of all metagenomic reads were assigned to 887 ARGs with a collective antibiotic resistance abundance index (ARAI)^20^ of 2.53 ppm. This included all major resistance mechanisms (Figure 3a): antibiotic target protection with 19 ARGs and 0.04 ppm (1.5%), antibiotic target replacement with 26 ARGs and 0.07 ppm (2.8%), antibiotic target alteration with 107 ARGs and 0.22 ppm (8.8%), antibiotic inactivation with 515 ARGs and 0.11 ppm (4.5%) and efflux-mediated resistance with 220 ARGs to an extraordinarily high share of collectively 2.07 ppm (82.4%).

**Figure 3:**
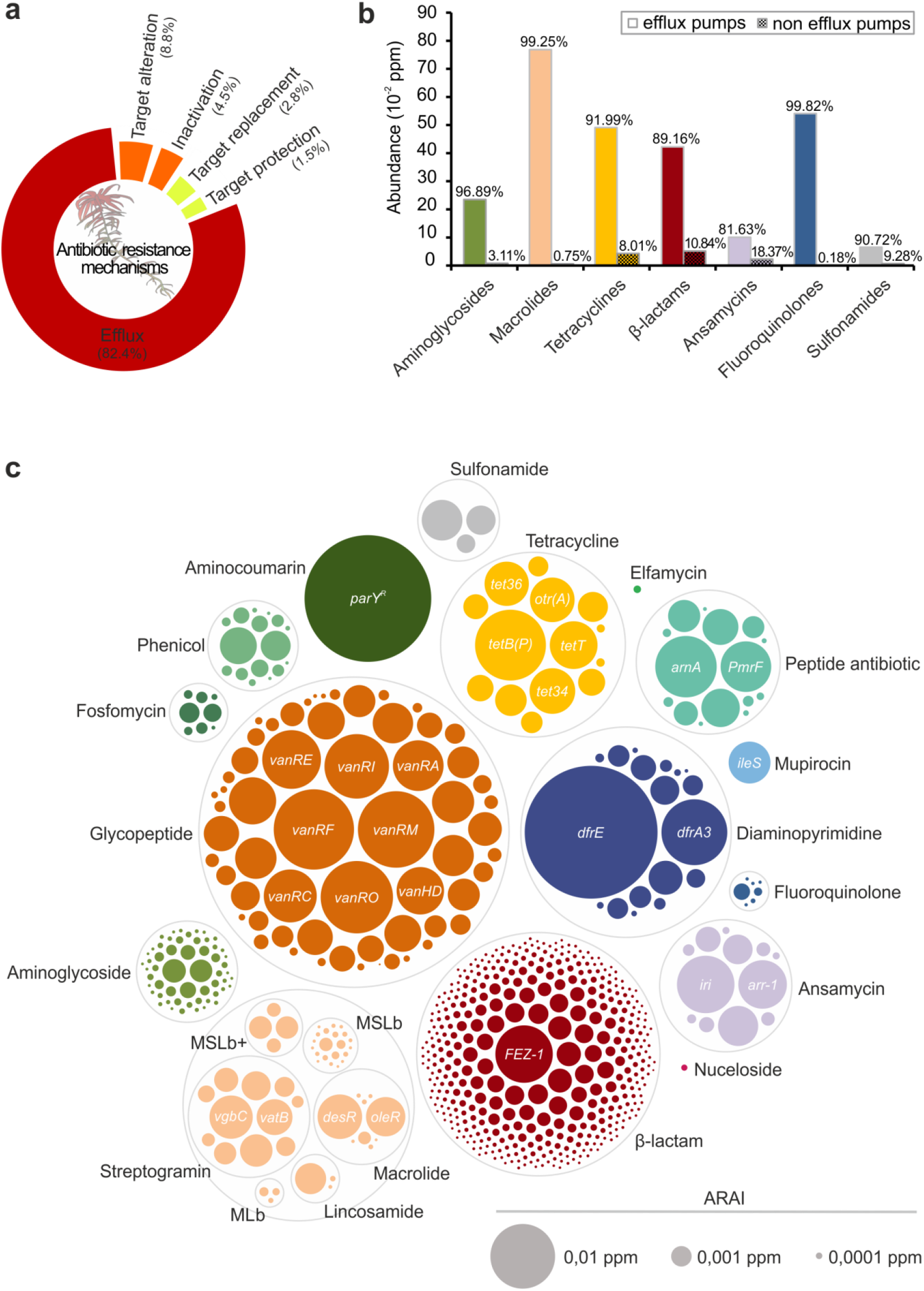
The *S. magellanicum* metagenome comprises a highly versatile resistome. The lllumina generated 41.8 Gbps moss metagenome was aligned against the CARD sequences. a) The five major resistance mechanisms presented by their relative abundance within the moss resistome. b) For a selected group of antibiotic classes, the extent of efflux pump mediated and non-efflux pump mediated resistance is compared. Abundance within the metagenome is given in absolute numbers by the ARAI in ppm (≙reads per million reads), while the abundance within antibiotic classes is given as proportion in percent. c) All detected non-efflux pump related ARGs grouped according to antibiotic classes. Each bubble represents one determinant with absolute abundance within the metagenome reflected by bubble size. The most abundant determinants are labelled with the gene names. MLb, macrolide and streptogramin. MSLb, MLb and streptogramin. MSLb+, MSLb and oxazolidione, pleuromutilin and phenicol. CARD, Comprehensive Antibiotic Resistance Database. ARAI, Antibiotic Resistance Abundance Index.

To understand the strong contribution of efflux pumps towards resistance in more detail, we evaluated the extent of this resistance mechanism against antibiotics at class level (Figure 3b). The focus was, thereby, restricted to the antibiotic classes used during the screening of the culture collection. Clycopeptides were omitted as these act on the outer cell wall^9^. Efflux pumps, which export multiple antibiotics, were included in the abundance of each of the respective antibiotic classes. Although to a varying degree between 80% to almost 100%, efflux pumps constituted the most abundant resistance mechanism for all studied classes. Efflux-mediated resistance is more prevalent for macrolides, tetracyclines, β-lactams and fluoroquinolones than for aminoglycosides, rifamycins and sulphonamides based on the determined ARAI.

Next, the detected resistance determinants were grouped according to their antibiotic class to compare for all classes their distribution and abundance. However, efflux pumps were excluded entirely in this analysis as they often confer resistance to multiple antibiotics. The overall target spectrum of the detected 667 non-efflux pump determinants spans 21 antibiotic classes including synthetic antibiotics such as diaminopyrimidines, fluoroquinolones and sulphonamides and many classes ranked as critically important for human medicine^18^ like aminoglycosides, glycopeptides and β-lactams (Figure 3c). These results show a high degree of genetic diversity and an even distribution of the detected ARGs as expected, ranging from 8.3 × 10^−6^ to 1.5 × 10^−2^ ppm (Figure 3c, Supplementary Data 2). Only two ARGs, *dfrE* and *parY*^*R*^ mark an exception being the most prominent determinants with a considerable difference in abundance with 4.1 × 10^−2^ and 3.8 × 10^−2^ ppm, respectively. β-Lactams represent the most abundant class with more than 400 ARGs. In contrast, for aminocoumarins, mupirocin, nucleoside and elfamycin just one ARG was assigned to each (*parY*^*R*^, *ileS, tmrB*, and *EF-Tu*). Almost all genetic determinants that were suggested by Berendonk *et al.*^8^ as potential indicators to survey the antibiotic resistance status in environmental samples were detected in this metagenome as well, although at lower abundance. These include *sul1, sul2, bla*_CTX-M_, *bla*_TEM_, *bla*_VIM_, *bla*_KPC_, *qnrS, vanA, mecA, ermB, ermF, tetM* and *aph* (Supplementary Data 2). Altogether, the data highlight the predominance of glycopeptide and β-lactam resistance determinants in the studied resistome, both in terms of abundance and versatility with 60 and 403 ARGs and 0.14ppm (32.6%) and 0.05 ppm (11.6%), respectively.

*In silico* analysis showed the predominance of β-lactam resistance determinants, and that β-lactamase diversity in the *Sphagnum* resistome covers every β-lactam class. The list of assigned β-lactamases included extended spectrum as well as metallo β-lactamases of environmental but also clinical origin, such as GIM-2, SHV-16 and TEM-102 (Supplementary Data 2). Due to the relevance of extended-spectrum and metallo β-lactamases, which pose a problem to the still widely administrated β-lactams, a network analysis was conducted to assess the target spectrum of the 398 assigned β-lactamases (Figure 4). All six β-lactam classes, penams, penems, monobactams, cephalosporins, cephamycins and carbapenems, are represented in the constructed network. The majority of determinants (67.6%) cluster in groups acting on more than one β-lactam class. These clusters often connect to three, four and five β-lactam classes comprising 22.1%, 13.3% and 6.5% of the detected determinants, respectively. Overall, the determinants connect most frequently to cephalosporins and penams. However, connections to carbapenems, drugs of last resort, are also highly represented in the network analysis.

**Figure 4:**
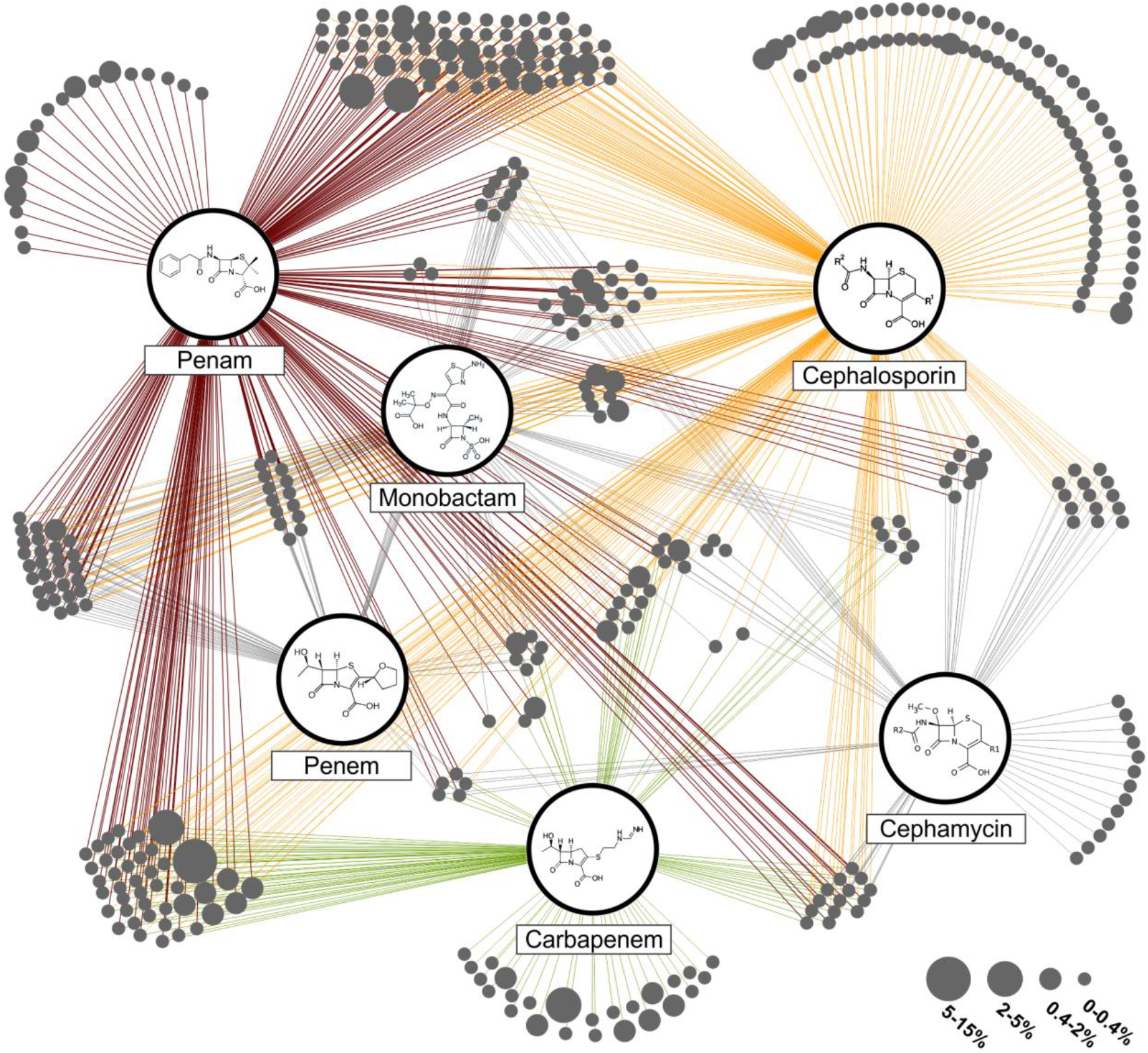
Substrate spectrum of detected β-lactamases. All assigned β-lactamases of the *S. magellanicum* metagenome represented as single bubbles (in dark grey) were grouped into clusters based on their reported substrate spectrum. Enzymes with the same substrate spectrum form one cluster. Connecting lines from the clusters to the β-lactam classes display the substrate specificity. Bubble size relates to the relative abundance of single enzymes within the whole β-lactamase pool. The three most abundant classes are penam (red), cephalosporin (orange) and carbapenem (green).

### Identification of the novel class A β-lactamase Mm3 from the *Sphagnum* metagenome

A functional metagenomics approach was pursued to identify novel resistance genes. Therefore, a fosmid library comprising 3.6 Gbps of cloned moss metagenomic DNA was screened for ARGs against nine different antibiotics. The screening identified three unique resistant metagenomic clones (*E*. *coli* EPI300 pCC2FOS-Mm1, Mm2 and Mm3), all three conferring resistance against ampicillin. The initially determined minimal inhibitory concentrations (MICs) for ampicillin were 64 μg ml^−1^ for clones Mm1 and Mm2, and >512 μg ml^−1^ for Mm3, as compared to 32 μg ml^−1^ for the control strain (Supplementary Table 2). The clone *E. coli* EPI300 pCC2FOS-Mm3, exhibiting the highest MIC for ampicillin, was chosen for *de novo* sequencing. This revealed a novel β-lactamase gene encoding a 304 amino acid protein with an estimated weight of 32.8 kDa to be present on the 40.7 kb DNA insert. The gene was designated *blaMm3* (<u>ß-la</u>ctamase from <u>M</u>oss <u>m</u>etagenome clone <u>3</u>).

The novel β-lactamase gene *blaMm3* shares the highest sequence similarity with two annotated but not yet characterised β-lactamases from *Rhodanobacter* sp. (70.6%) and *Frateuria* sp. (66.8%). Both species belong to the family of *Rhodanobacteraceae* and order of *Xanthomonadales*. Together with reference sequences from characterised β-lactamases the phylogenetic relation of Mm3 was elucidated (Figure 5). Its clustering into defined groups of class A β-lactamases was evaluated according to the updated classification by Philippon *et al*.^21^. The Mm3 β-lactamase clustered, together with the next neighbour sequences from *Rhodanobacter* sp. and *Frateuria* sp., in closer proximity to members of the so called *Xanthomonas* (XANT) group, which contains β-lactamases from *Xanthomonas* sp., *Stenotrophomonas maltophilia* and *Pseudomonas aeruginosa*. Other clusters in the phylogenetic tree include members showing a lower degree of similarity (33 to 44% identity), like those belonging to the limited-spectrum (LSBL1 to 4) and extended-spectrum β-lactamases (ESBL1 and 3). Members of the LSBL2 and 3 clusters have been described as true carbenicillinases, while enzymes from the ESBL group hydrolyse cephalosporins like cefotaxime additionally to penicillins. In accordance with the phylogenetic analysis, the amino acid sequence of Mm3 harbours characteristic class A Ambler motifs^22^ as follows: 70SerThrPheLys (SxxK motif), 130SerAspAsn (SDN motif), 234LysThrCly (KTC motif), Glul66 and 166GluProGluLeuAsn (ExxLN motif).

**Figure 5:**
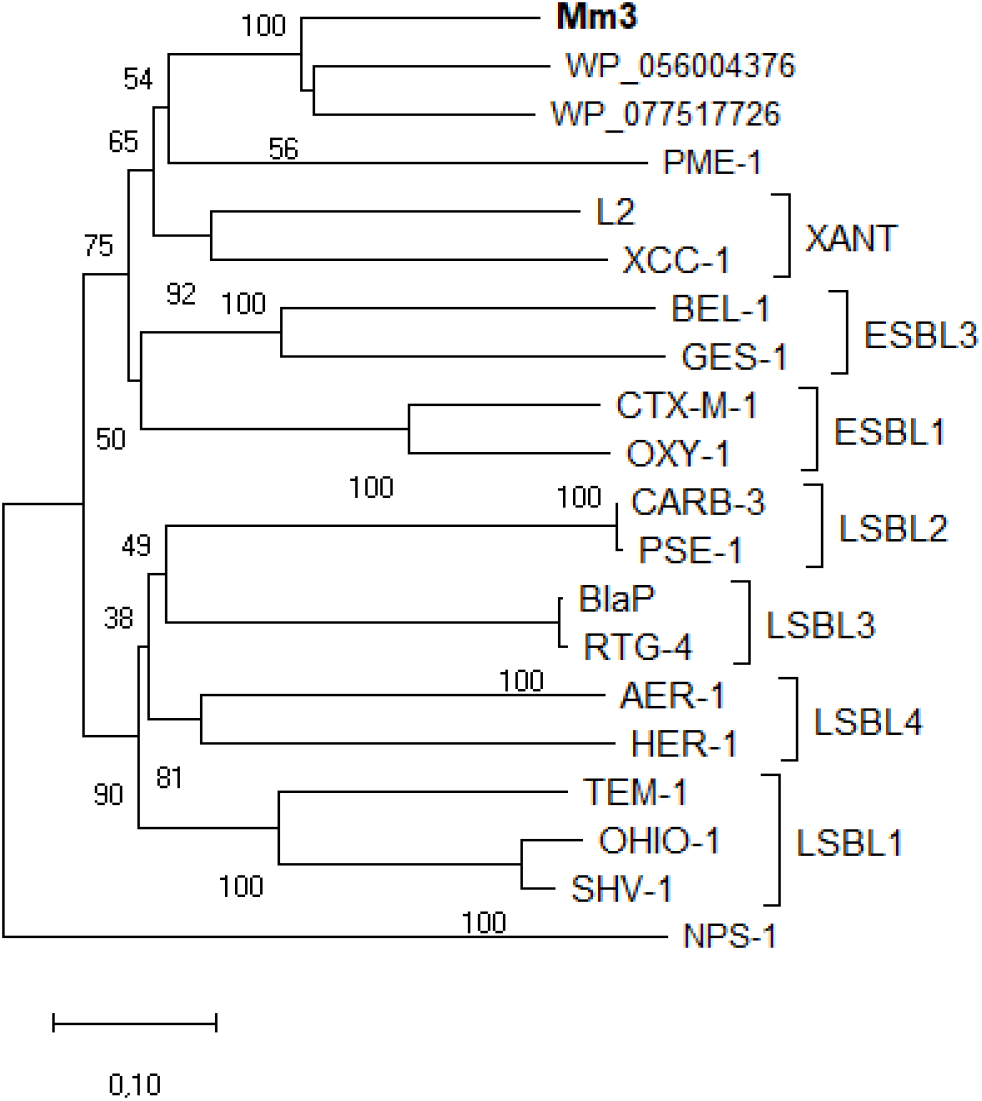
Phylogenetic relationship of Mm3 and other class A β-lactamases. The evolutionary analysis of aligned amino acid sequences was conducted using the neighbor-joining method. Bootstrap values are shown next to the branches. The scale bar indicates the number of amino acid differences per sequence. The reference sequences are: *Frateuria* sp. Soi1773 (WP_056004376), *Rhodanobacter* sp. C03 (WP_077517726), PME-1 (*Pseudomonas aeruginosa*, E9N9H5), L2 (*Stenotrophomonas maltophilia*, P96465), XCC-1 (*Xanthomonas campestris* pv. *campestris*, 087643), BEL-1 (*Pseudomonas aeruginosa*, Q3SAW3), GES-1 (*Klebsiella pneumonia*, Q9KJY7), CTX-M-1 (*Escherichia coli*, P28585), OXY-1 (*Klebsiella oxytoca*, P22391), CARB-3 (*Pseudomonas aeruginosa*, P37322), PSE-1 (*Pseudomonas aeruginosa*, Q03170), AER-1 (*Aeromonas hydrophila*, Q44056), HER-1 (*Escherichia hermannii*, Q93FN7), BlaP (*Proteus mirabilis*, P30897), RTG-4 (*Acinetobacter baumannii*, ACJ61335), TEM-1 (*Shigella flexneri*, AAC97980), OHIO-1 (*Enterobacter cloacae*, P18251), SHV-1 (*Klebsiella pneumonia*, P0AD64). The tree was rooted with NPS-1, a class D β-lactamase from *Pseudomonas aeruginosa* (AAK1479).

To investigate the substrate spectrum of the novel β-lactamase, MICs for penam and cephalosporin antibiotics were determined (Supplementary Table 2). Similarly to the control strain, clone Mm3 showed no to little resistance to the tested cephalosporin concentrations with MICs of <0.5, 64 and 8 μg ml ^−1^ for cefatoxime, cephalexin and cefalothin, respectively. On the contrary, Mm3 displayed a distinctly higher resistance against the penam antibiotics ampicillin (>512 μg ml^−1^) and carbenicillin (>1024 μg ml^−1^).

The identified *blaMm3* gene encoding a novel β-lactamase was cloned and expressed in *E. coli* BL21 (DE3) for subsequent purification and biochemical characterisation. After confirming solubility of the overexpressed enzyme by SDS-PACE analysis, β-lactam-hydrolysing activity was verified by testing cell-free lysates on nitrocefin disks (data not shown). The N-terminally His-tagged enzyme was then purified by affinity chromatography to a purity of 90% as estimated by SDS-PAGE. Two prominent bands with a molecular weight of around 32 and 35 kDa were visible (Supplementary Figure 1). LC-MS/MS analysis of both bands determined each of the respective proteins to comprise the right β-lactamase amino acid sequence (data not shown). Flowever, the 35 kDa protein contained the N-terminal Flis-Tag while the lower weight protein did not (32 kDa), probably as a result from proteolytic activity during purification or SDS-PAGE analysis. Data obtained from the kinetic measurements of Mm3 for ampicillin were fitted using the Hill equation (n_H_ of 2.36). In the case of carbenicillin the kinetic data were fitted with the Michaelis-Menten equation (Supplementary Figure 2). The kinetic analysis revealed a higher affinity of Mm3 for ampicillin (V_max_ = 179.2 ± 6.1 U mg^−1^, K_M_ = 270.8 ± 16.4 μM) compared to carbenicillin (V_max_ = 264.6 + 8.6 U mg^−1^, K_M_ = 399.85 ± 42.69 μM).

## DISCUSSION

Our multi-faceted analysis uncovered a highly versatile resistome present in the evolutionary old and long-term stable bog ecosystem, and underlined the natural, strong resilience of *Sphagnum*-associated bacteria against antibiotics. Given the highly adapted plant-associated lifestyle, the strong microbial competition and the vast pool of microbial and plant-produced secondary metabolites^12^, the *Sphagnum* microbiota has developed general and also specific antimicrobial resistance mechanisms that naturally equip them against antibiotics. (Semi)synthetic drugs were not exempt from this as demonstrated in the present study. Contrary to our initial expectation, a predominance of resistances against synthetic antibiotics was observed in the culture collection, despite its pristine origin^10^. Stemming to a large part from efflux pumps as indicated by our *in silico* analysis, the ability to combat these compounds may not exclusively result from extrusion. For instance, resistance of environmental bacteria against synthetics has been tied to high sequence variations in target genes^23^. Furthermore, a high level of multi-resistances was encountered in the culture-dependent analysis. The six isolates with the highest level of multi-resistance belonged to *Burkholderiales* and *Enterobacteriales*, which are typical and dominant orders within the bacterial community of *Sphagnum* mosses^12,14^. *S. marcescens, P. phytofirmans*, and *R. chamberiensis* were isolated from this habitat before^16, 24, 25^. For *Pandoraea* spp. an association with *Sphagnum* was not described so far. Interestingly, this bacterium has been mostly isolated from the sputum of cystic fibrosis patients^26^ and is considered as emerging opportunistic pathogen^27^. Clinically isolated *Pandoraea* spp. are known to be highly resistant, including resistances to last defence antibiotics of the carbapenem class^27, 28^. The two multi-resistant isolates described here show that these species naturally possess a high level of resistance and they might easily transit to clinical environments as they are well equipped with ARGs. *S. marcescens*, a common plant-associated bacterium, also exhibits various antibiotic resistances and opportunistic traits^29^. Less is known to this end for the identified *Paraburkholderia* and *Rouxiella* species, especially for *P. phytofirmans*, which is a promising plant growth promoting agent^30^. Interestingly, plants in general and *Sphagnum* in particular constitute reservoirs for plant growth promoting bacteria with antifungal and antibacterial activity^31^, while simultaneously hosting species known as opportunistic human pathogens^16,32^. According to our results, assessment of the environmental resistome in a given habitat can be used to predict emerging opportunistic pathogens and thereby help to counteract bacterial infections, which are considered a serious public health issue world-wide^33^.

The high microbial diversity of the *Sphagnum* moss microbiome was reflected by the corresponding resistome, which displayed high versatility and evenness. Given the high stringency applied to our analysis, the observed coverage on the functional and chemical level with more than 800 ARG-like genes covering 21 antibiotic classes is staggering. A key driver for this vast diversity resides in the presence of a great repertoire of efflux pumps. They are considered as an evolutionary ancient and general resistance mechanism against toxic molecules like heavy metals, solvents, and plant-produced antimicrobials^34^, and may be a missing link in understanding resistome composition in natural environments. For the *Sphagnum* resistome efflux-mediated resistance comprised more than 80%. An extraordinarily high share when compared to other CARD-based analyses, which attributed a relative proportion of 20-50% of the resistances to efflux pumps for various natural and human controlled environments^6,35,36^. Yet, the high share of efflux pumps is a common attribute for other plant-associated resistomes in *Sphagnum*-dominated peatlands (unpublished data). As their typically high taxonomic diversity is inherently a driving force for chemical diversity, efflux pumps, thus, constitute a pivotal point in ensuring co-existence within this highly complex community. This general resistance mechanism fosters the diverse pool of ARGs present in the moss resistome and in doing so contributes to the great plasticity found within the peat bog ecosystem.

Since *Sphagnum* mosses are rootless plants that within *Sphagnum*-dominated peat bogs do not have soil contact, the elucidated resistome is regarded as intrinsic. As such, it addresses the need to understand the extent to which the plant resistome is intrinsic or recruited from soil, which represents still an unanswered question^5^. In addition, we expect that it will be vertically transmitted with the core microbiome from the gametophyte to the sporophyte and *vice versa*^14^. Our data highlight that the plant microbiome naturally comprises a versatile, intrinsic resistome. This is reinforced by the identification of a novel class A β-lactamase. Notably, Mm3 shares low sequence similarity to characterised β-lactamases. This can be explained by the fact that most moos-associated microorganisms are not yet cultivable and not much is known about their origin and genetic content^12^. The metagenome derived β-lactamase Mm3 is phylogenetically closest related to β-lactamases of the XANT group and the uncharacterised β-lactamases from the environmental isolates *Rhodanobacter* (soil) and *Frauteuria* (rhizosphere), both belonging to the order of *Xanthomonadales*^37, 38^, *Xanthomonadales spp*. constitute common colonisers of *Sphagnum* mosses^39,40^. Interestingly, many of the other β-lactamase sequences clustering in close proximity were isolated from well-known nosocomial human pathogens such as *S. maltophilia, P. aeruginosa* or *K. pneumoniae*^27^. The relatedness of *bla*Mm3 to these genes is not surprising, since β-lactamases account as evolutionary old enzymes and are widely spread in nature^9^. The latter was clearly confirmed in the network analysis, displaying high abundance and an extraordinary diverse substrate range of the *in silico* detected β-lactamases. The isolated β-lactamase Mm3 showed a higher affinity for penam antibiotics, but no activity for the tested cephalosporins, exhibiting in this case a narrow substrate spectrum. With K_M_ values around 270 to 400 μM for the penam antibiotics, the activity of the new Mm3 is not outstanding and surpassed by the ones reported for β-lactamases from many facultative human pathogens, e.g. the plasmid-encoded MIR-1, CMY-1 or ACT-1 from *E. coli* (0.16 to 2.2 μM, ampicillin)^41–43^. Based on this finding, the presence of antibiotic-inactivating enzymes with a rather limited activity seems to be characteristic for the described natural environment as compared to the clinical settings.

Since ARGs are often associated with specific taxa^36,44,45^, we expected the taxonomically diverse and balanced *Sphagnum*-associated microbiome to comprise a resistome with evenly distributed ARGs at low abundance; a common observation for natural environments^4^. In contrast, environments under anthropogenic influence are characterised by highly abundant ARGs^4,36^, and further correlate with a loss in bacterial diversity and enrichment of opportunistic pathogens^36^. The antimicrobial selective pressure exerted by our life style is without question the driving force for imbalance, leading to a shift in bacterial community composition that ensues the increase of opportunistic pathogens and their associated ARGs. However, overlooked in this context is the ecological concept of *K-* and *r*-selection favouring oligotrophic or copiotrophic taxa, respectively. We recently reported that the phyllosphere of arugula from urban gardening was dominated by *Gammaproteobacteria* and in particular by multi-resistant *Enterobacteriacaea*^6^. This class, which comprises many opportunistic pathogens, tends towards a copiotrophic lifestyle, displaying faster growth and substrate generalisation as compared to the more oligotrophic *Alphaproteobacteria*^46^. The *Sphagnum* microbiota in contrary is dominated by *Alphaproteobacteria*, such as the slow growing *Methylobacteria*^12,14^. We assume that *K*-selection maintains oligotrophy and stabilises the bacterial community in the nutrient poor, microbial rich ecosystem of *Sphagnum-*dominated bogs. In doing so it represents a driving force in shaping the observed evenness and diversity within the moss resistome. Microbial community management ensuing diverse, stable and beneficially designed microbiomes is foreseen to abate exposure to resistances^36^. We propose that resistance management in form of microbial community management could be achieved through *K*-selection. The advantageous effects of such a strategy have already proven valuable in improving the larvae viability in aquaculture^47^.

Based on our complementary screening strategy, the herein presented novel findings deliver a first comprehensive picture of a native plant resistome consisting of a highly diverse genetic pool and novel antibiotic resistance genes.

## METHODS

### CARD-based *Sphagnum* resistome profiling

A 41.8 Gbps metagenome previously generated by lllumina HiSeq paired-end sequencing from the Alpine peat bog moss *S. magellanicum*^12^ was implemented for antibiotic resistance profiling. The 172 590 841 paired-end reads were aligned against sequences from the CARD^48^, retrieved in April 2017, using the diamond protein aligner v0.9.24^49^. BLASTX^50^ was performed at high stringency with a similarity threshold of 90% over the full read length and otherwise default settings giving hits with ≥30 amino acids identity. The reads were assigned to their best BLASTX hit and the obtained dataset was manually curated for gene redundancy or in case of antibiotic target genes for known resistance conferring mutations (Supplementary Table 1). The reads were normalised by calculating the ARAI (number of reads assigned to one ARG per number of total reads and respective ARG length)^20^. Abundance of ARGs or resistance mechanism within the metagenome is given as ppm (≙read per million reads), while their proportion among all assigned ARGs or resistance mechanisms is given as percentage. The detected non-efflux pump determinants were visualised using RAWGraphs^51^. The distribution and abundance network of assigned β-lactamases was constructed with Cytoscape v3.3.0^52^.

### Sampling and isolation of *S. magellanicum* associated bacteria

Gametophyte samples of the moss *S. magellanicum* were collected from the Austrian Alpine bog Pürgschachen Moor (N47°34′50.57″ E14°20′29.29″) in September 2017 (Figure 1). Fluorescent *in situ* hybridization and confocal laser scanning microscopy were performed on *Sphagnum* gametophytes as described previously using the reported probes (Cy3-labeled ALF968 for *Alphaproteobacteria*, Cy5-labeled EUB338, EUB338II and EUB338III for *Eubacteria*)^53^.

The cleaned and fractionated plant material was shaken in a Stomacher laboratory blender (BagMixer, Interscience, France) twice for 120 s batch-wise in sterile plastic bags to 20 g in 50 ml of chilled 0.85% NaCI solution. After straining the suspension through double-layered gauze and a sterile analysis sieve (mesh size 63 pm, Retsch, Germany), the undiluted suspension as well as serial dilutions thereof were plated on R2A agar (Roth, Germany) containing nystatin (25 μg ml^−1^, Duchefa Biochemie, Netherlands) and incubated at 20°C for four days. Isolates were subcultured until purity and liquid cultures grown from single, isolated colonies in Nutrient Broth II (Sifin diagnostics, Germany) supplied with glycerol to 20% (v/v) for long term storage at −70°C.

### Antibiotic resistance screening of the *Sphagnum* culture collection

Bacterial isolates were screened against ten different antibiotics as listed in Supplementary Table 3. Concentrations were based on those used in other studies^6,54^, which followed the guidelines by the Clinical Laboratory Standards Institute. All 264 isolates were transferred to R2A agar plates of up to 50 isolates per plate and incubated at 20°C for four days. The colonies were then replica printed onto Müller-Hinton agar plates supplemented with the different antibiotics and incubated at 20°C. The plates were monitored every 24 hours for three days. This was done in duplicate and only isolates which had grown to visible colonies in both screenings were considered resistant.

### Annotation of multi-resistant isolates

Cells were mechanically disrupted by bead-beating (6 m s^−1^, 20 s) and the lysates incubated at 95°C for 10 minutes followed by centrifugation at 5000 rpm for 5 min. The 16S rRNA gene was amplified using 2 μl of the supernatant and the universal bacterial primer pair 27f (5’AGAGTTTGATCMTGGCTCAG) and 1492r (5’TACGGYTACCTTGTTACGACTT), 0.5 μM each, in a 50 μl PCR reaction with lx Taq-&GO Ready-to-use PCR Mix (MP Biomedicals, Germany) (98°C – 4 min; 25 cycles of 98°C – 30 s, 48°C – 30 s, 72°C – 90 s; 72°C – 5 min). The 1400 bp long DNA fragments were purified (Wizard^®^SV Gel and PCR Clean-Up System, Promega, Germany) and sequenced using the same 27f and 1492r primer pair. Sequences were annotated to their best NCBI hit.

### Metagenomic library construction

The 3.6 Gbps metagenomic fosmid library was established in *E. coli* EPI300 pCC2FOS (Epicentre, Wisconsin, USA), incorporating ~40 kb metagenomic DNA from the Alpine peat bog moss *S. magellanicum* as previously described^55^. The generated 90 000 metagenomic clones were pooled by resuspending them in LB medium supplied with 20% glycerol for long term storage at −70°C.

### Screening for novel antibiotic resistance genes

The functional screening of the *Sphagnum* metagenomic library was carried out on LB agar plates containing chloramphenicol (12.5 μg ml^−1^) for fosmid maintenance and arabinose (0.01% w/v) to induce high-copy number. Metagenomic clones were screened against nine different antibiotics as listed in Supplementary Table 3. Concentrations were chosen according to those used in other studies employing the CopyControl system with *E. coli* EPI300 ^56,57^. Alternatively, the MIC was determined as described below. Cells of the pooled library stock were revived in LB broth containing chloramphenicol (12.5 μg ml ^−1^) at 37°C for 3 h with shaking at 130 rpm. The library was screened with at least 3x coverage by plating 50 000 to 100 000 CFU per plate. Colonies that had formed after 16 h of incubation at 37°C were re-cultivated under the same conditions to confirm the phenotype. Resistant clones were evaluated by restriction digest and unique clones were retransformed to confirm the presence of the resistance phenotype on the fosmid insert.

### *De novo* sequencing of pCC2FOS-AmpR3

Extracted DNA was used to generate Illumina shotgun paired-end sequencing libraries, which were sequenced with a MiSeq instrument and the MiSeq reagent kit version 3, as recommended by the manufacturer (lllumina, California, USA). Quality filtering using Trimmomatic version 0.36^58^ resulted in paired-end reads with an average read length of 301 bp. The assembly was performed with the SPAdes genome assembler software version 3.10.0^59^, resulting in a 50.2 kb contig with a 9.2-fold coverage. The assembly was validated, and the read coverage determined with QualiMap v2.1^60^. Automatic gene prediction was performed using the software tool Prokka v1.12^61^.

### Phylogenetic analysis of *blaMm3*

The phylogenetic analysis for inferring the evolutionary relationship of Mm3 with other β-lactamases was conducted using the software MEGA X v10.0.2^62^. The amino acid sequences were aligned using MUSCLE^63^ and the tree was constructed by the neighbour-joining method^64^ with a bootstrap test of 2000 replicates, using the p-distance method^65^ for computing evolutionary distances.

### Minimum inhibitory concentrations

MICs were determined according to the guidelines of the European Committee for Antimicrobial Susceptibility Testing using the broth microdilution method^66^. The assays were conducted in triplicate. MICs for the functional metagenomics screening were determined using the empty vector library host *E. coli* EPI300 pCC2FOS. MICs for the ampicillin resistant clones were determined for ampicillin and for *E. coli* EPI300 pCC2FOS-Mm3 additionally for cefotaxime, cephalothin, cephalexin, carbenicillin (Sigma-Aldrich, Germany) using *E. coli* EPI300 pCC2FOS as control strain.

### Subcloning *blaMm3*

The *blaMm3* gene was cloned into the pET28a(+) expression vector (Novagen, USA) with N-terminal His Tag and inducible T7 promoter using the Ndel and EcoRI restriction sites. With primers comprising the respective restrictions sites (underlined) (F: 5’-3’ TGCAGA<u>CATATG</u>AACCCCAACCACTCTG. R: 5’-3’ TACT<u>AGAATTC</u>CTAGACGCTCGATGTCGCC, Sigma-Aldrich, Germany), the full ORF was amplified from pCC2FOS-Mm3 by a standard PCR reaction using the Phusion DNA polymerase (New England BioLabs, Germany) at 72°C annealing temperature. The vector ligated gene was transformed into high efficiency *E. coli* DH5α (New England BioLabs) for selection of the recombinant pET28a*-blaMm3* plasmid, which was then introduced into *E. coli* BL21(DE3) (Thermo Scientific, Germany) for overexpression.

### Expression and purification of the β-lactamase Mm3

LB broth (400 ml) with kanamycin (50 μg ml^−1^) was inoculated with 2% (v/v) of an overnight culture of *E*. *coli* BL21(DE3) pET28a*-blaMm3* which was then grown at 37°C under shaking at 130 rpm to an OD_600_ of 0.8. The culture was supplemented with isopropyl β-D-1-thiogalactopyranoside to 0.4 mM end concentration and further incubated for 4 hours. Harvested cells were resuspended in 50 ml binding buffer (20 mM sodium phosphate buffer, 500 mM NaCI, 20 mM imidazole, pH 7.4) containing 0.8 g I^−1^ lysozyme and disrupted by sonication with a digital sonifier (pulses of 2 s and 4 s pause, 5 min, *70%* amplitude; Branson, Emerson, Missouri, USA). The His-tagged protein was isolated from the centrifuged lysate (12000 × g, 10 min) using a 1-ml HisTrap column (GE health Care, Illinois, USA) and an elution gradient (1 ml min^−1^, 20 min) up to 500 mM imidazole. Two fractions containing active β-lactamase, as judged by application on nitrocefin disks (Sigma Aldrich, Germany) and SDS-PAGE were mixed together. The buffer was exchanged with 20 mM sodium phosphate buffer (200 mM NaCI, pH 7.4) through multiple dilution and centrifugation steps using Amicon Ultra-15 centrifugal filters (10 kDa cuttoff, Merck Millipore, Germany). Protein purity was estimated with SDS-PAGE and the concentration determined with the Pierce BCA assay kit (Thermo Scientific) using bovine serum albumin as reference. The purified enzyme was shock-frozen in liquid nitrogen and stored at −70°C.

### Kinetic characterisation of the ß-lactamase Mm3

To determine kinetic values (V_max_ and K_m_) the activity of the purified β-lactamase Mm3 was measured spectrophotometrically (U-2001, Hitachi, Tokyo, Japan). Initial hydrolysis rates for ampicillin and carbenicillin were recorded at 235 nm and 30°C in 450 μl reaction buffer (20 mM sodium phosphate buffer, 0.2 M NaCI, pH 7.4) upon addition of 50 μl substrate at different concentrations (1 mM to 100 mM in H_2_O). The kinetic data was fitted for ampicillin with the Hill and for carbenicillin with the Michaelis-Menten equation, respectively, using the software Origin 9.0.0G (Origin Lab Corporation, Massachusetts, USA).

### Data accessibility

The complete *S. magellanicum* metagenome is stored at the MGRAST server under the accession no. 4533611.3. The nucleotide sequence of the β-lactamase *blaMm3* is deposited in Genbank under the accession no. MK831000 and the 16S rRNA sequences from the multi-resistant bacteria under the accession numbers MK801238-MK801243.

## Supporting information

Supplementary material: Data 1 Isolate resistance profiles

Supplementary material: Data 2 CARD data

## ACKNOWLEDGMENTS

We thank Christian Berg for sampling of *S. magellanicum*, and Stephanie Hollauf, Franz Stocker and Angelika Schäfer for helping with the preparation of moss samples for downstream processing, Harald Blasl and Bettina Semler for their help with the screenings and Silvia Ferrario for helping with subcloning and protein expression. Special thanks go to Isabella Wrolli for her indispensable technical help throughout the entire study as well as to Henry Müller (all Graz) for his technical guidance. This work was supported by the Federal Ministry of Science, Research and Economy (BMWFW), the Federal Ministry of Traffic, Innovation and Technology (bmvit), the Styrian Business Promotion Agency SFG, the Standortagentur Tirol, the Government of Lower Austria and ZIT – Technology Agency of the City of Vienna through the COMET-Funding Program managed by the Austrian Research Promotion Agency FFG.

## CONTRIBUTIONS

M.-M.O., C.A.M. and G.B. conceived the study. M.-M.O. and C.A.M. designed and performed the experiments. M.-M.O, J.T., T.C. and A.B. analysed and/or interpreted the data. A.P. performed *de novo* sequencing and annotation. M.-M.O., C.A.M., and G.B. wrote the paper. All authors revised the manuscript and approved the final version.

