## Supplementary material: Figures and Tables for "Unravelling native plant resistomes – The *Sphagnum* microbiome harbours versatile and novel antimicrobial resistance genes"

Melanie-Maria Obermeier[1,2], Julian Taffner[1], Alessandro Bergna[1,2], Anja Poehlein[3], Tomislav Cernava[1], Christina Andrea Müller[1,2]*, Gabriele Berg[1,2]

[1]Institute of Environmental Biotechnology, Graz University of Technology, Graz, Austria
[2]Austrian Centre of Industrial Biotechnology, Graz, Austria
[3]Goettingen Genomics Laboratory, Georg-August-University Goettingen, Goettingen, Germany

*Corresponding author:

Christina Andrea Müller
Institute of Environmental Biotechnology
Graz University of Technology, Petersgasse 12
8010 Graz, Austria


### FIGURES

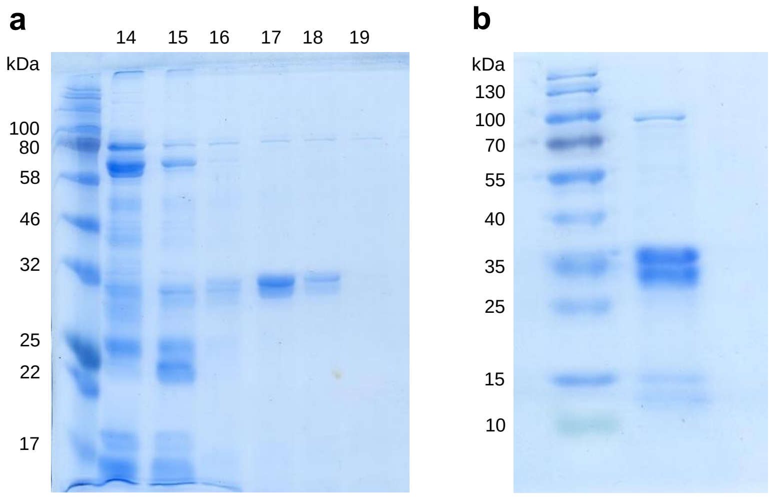


**Supplementary Figure 1: Purified β-lactamase Mm3.** The *blaMm3* gene encoding a 304 amino acid long β-lactamase was cloned under an N-terminal His Tag and the tagged protein purified by affinity chromatography. a) Elution fractions no. 14 to 19; no. 17 and 18 were selected for further use. b) SDS-PAGE of the pure enzyme shows two bands, one with the estimated molecular weight of the his-tagged protein of about 35 kDa and a smaller protein band around 32 kDa. As identified by LC-MS/MS analysis the higher molecular weight band contains the β-lactamase still adjunct to the His-Tag, while the lower band stems from a smaller version of the purified β-lactamase which lost the His-Tag. Lower molecular weight proteolytic products of approximately 15 to 17 kDa are visible as well, corresponding to the His-tagged termini of the protein. Protein Ladder: a) Color prestained Protein Standard, Broad Range (New England Biolabs); b) PageRuler Prestained Protein Ladder (Thermo Scientific).

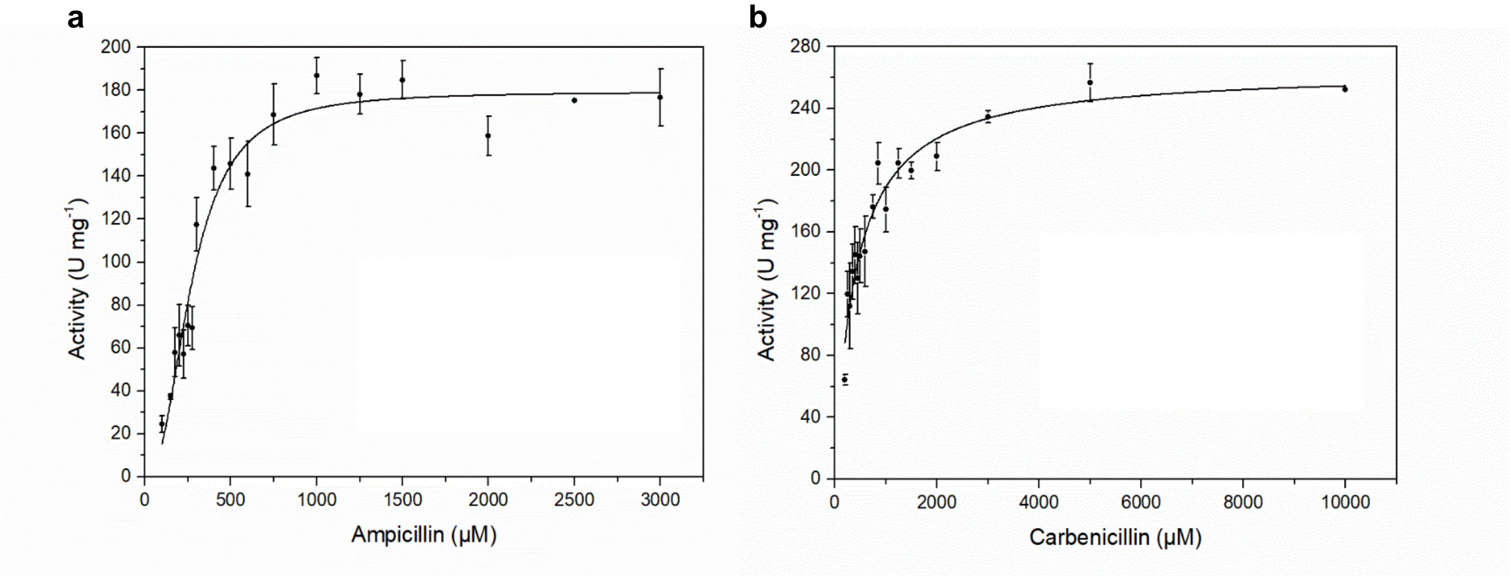


**Supplementary Figure 2: Kinetic characterisation of the β-lactamase Mm3.** The initial hydrolysis of the substrate was followed spectrophotometrically at 235 nm. Points, mean values (n = 2 to 8) for all measurements, except for 2.5 mM carbenicillin (n=1); error bars, standard deviations. Data for ampicillin (a) and carbenicillin (b) were fitted according to the Hill or Michaelis-Menten equation, respectively.

### TABLES

**Supplementary Table 1: List of manually curated resistance determinants.** Abundance is given in number of assigned reads before and after curation and after normalisation of the curated abundance as ARAI*. a) Genes which represent antibiotic targets, for which point mutations are known to confer resistance. Assigned reads which align to the mutation area were filtered and only those containing the point mutation were retained in the dataset. b) Genes and their assigned Antibiotic Resistance Ontologies (AROs) to which two or more gene sequences are attributed to. To avoid multiple representations of such AROs, hits of homolog genes were merged. c) Genes sourced out due to high similarity to common and widely spread genes, which are not directly related to antibiotic resistance.

| | Gene (ARO) | Abundance before curation | A) Mutation (No. of reads aligning to mutation area) B) Accession numbers of homologous genes | Abundance after curation | ARAI* (ppm) after normalisation of curated reads |
|---|---|---|---|---|---|
| A) Curated for mutations | *Chlamydia trachomatis murA* (ARO:3003785) | 194 | C119D (0) | 0 | 0 |
| | *Mycobacterium tuberculosis murA* (ARO:3003784) | 1058 | C117D (309) | 22 | 0.00030495 |
| | *Streptomyces cinnamoneus EF-Tu* (ARO:3003359) | 13978 | A379T (1039) | 9 | 0.00013135 |
| B) Curated for gene redundancy | *arnA* (ARO:3002985) | 361 | NP_252244 | 1007 | 0.00884032 |
| | | 646 | AAC75315.1 | | |
| | *cat* (ARO:3002670) | 114 | AAA22081.1 | 118 | 0.00317999 |
| | | 2 | AAA23018.1 | | |
| | | 1 | BAC11901.1 | | |
| | | 1 | AAB23649.1 | | |
| | *catIII* (ARO:3002685) | 1 | CAB75601.1 | 2 | 5.4404E-05 |
| | | 1 | CAA30695.1 | | |
| | *ANT(6)-Ib* (ARO:3002629) | 1 | CBH51824.1 | 2 | 4.0236E-05 |
| | | 1 | AIJ27543.1 | | |
| C) Other | *mfd* (ARO:3003844) | 8097 | ** | 0 | 0 |
| | *NmcR* (ARO:3003665) | 442 | *** | 0 | 0 |

*ARAI (antibiotic resistance abundance index): number of reads assigned to an antibiotic resistance gene per total number of reads and respective gene length in ppm (≙reads per million reads)[1].

** *Mfd* influences the spontaneous mutation rate that can give rise to ciprofloxacin resistances in *Campylobacter jejuni*[2]. It is, however, a wide-spread protein involved in DNA repair and by itself not directly related in antibiotic resistance[3].

*** *NmcR* regulates the *NmcA* β-lactamase, to which one metagenomic read was assigned resulting in an abundance of 1,98E-05 ppm (Supplementary Data 2). However, *NmcR* is a homolog of the widely conserved *lysR* regulators[4].

**Supplementary Table 2: Minimal inhibitory concentrations of β-lactam resistant metagenomic clones.** Minimal inhibitory concentration (µg ml$^{-1}$) of penam and cephalosporin antibiotics for the metagenomic clones *E. coli* EPI300 pCC2FOS-Mm1, Mm2 and Mm3 and the empty vector control *E. coli* EPI300 pCC2FOS (X). Mean values, n = 3.

| Clone | Ampicillin | Carbenicillin | Cefotaxime | Cefalothin | Cephalexin |
|---|---|---|---|---|---|
| Mm1 | 64 | 16 | 8 | 8 | 4 |
| Mm2 | 64 | 32 | 8 | 8 | 8 |
| Mm3 | >512 | >1024 | <0.5 | 64 | 8 |
| X | 32 | 8 | <0.5 | 8 | 4 |

**Supplementary Table 3: Antibiotic concentrations for resistance screenings used in this study.**

| Antibiotic | Antibiotic class | Manufacturer | Spectrum | Concentration [µg ml $^{-1}$] used for | |
|---|---|---|---|---|---|
| | | | | Isolates | Metagenomic clones |
| Ampicillin | β-Lactam | Roth, Germany | Gram +/- | 10 | 50 |
| Ciprofloxacin | Fluoroquinolone | Sigma-Aldrich, Missouri, USA | Gram +/- | 5 | 1 |
| Erythromycin | Macrolide | Roth, Germany | Gram +/- | 15 | 150 |
| Gentamycin | Aminoglycoside | Roth, Germany | Gram +/- | 10 | 10 |
| Kanamycin sulfate | | Roth, Germany | Gram +/- | 30 | 20 |
| Nalidixic acid | Quinolone | Merck, Germany | Gram +/- | 30 | 15 |
| Tetracycline | Tetracycline | Merck, Germany | Gram +/- | 30 | 4 |
| Rifampicin | Ansamycin | Duchefa Biochemie, Netherlands | Gram +/- | 5 | 20 |
| Sulfadiazine | Sulfonamide | Sigma-Aldrich, Missouri, USA | Gram +/- | 300 | ≤ 2250 |
| Vancomycin | Glycopeptide | Sigma-Aldrich, Missouri, USA | Gram + | 30 | 1000 |
